# Identification and characterization of putative *Aeromonas* spp. T3SS effectors

**DOI:** 10.1101/570887

**Authors:** Luiz Thiberio Rangel, Jeremiah Marden, Sophie Colston, João Carlos Setubal, Joerg Graf, J. Peter Gogarten

## Abstract

The genetic determinants of bacterial pathogenicity are highly variable between species and strains. However, a factor that is commonly associated with virulent Gram-negative bacteria, including many *Aeromonas* spp., is the type 3 secretion system (T3SS), which is used to inject effector proteins into target eukaryotic cells. In this study, we developed a bioinformatics pipeline to identify T3SS effector proteins, applied this approach to the genomes of 105 *Aeromonas* strains isolated from environmental, mutualistic, or pathogenic contexts and evaluated the cytotoxicity of the identified effectors through their heterologous expression in yeast. The developed pipeline uses a two-step approach, where candidate families are initially selected using HMM profiles with minimal similarity scores against the Virulence Factors DataBase (VFDB), followed by strict comparisons against positive and negative control datasets, greatly reducing the number of false positives. Using our approach, we identified 21 *Aeromonas* T3SS likely effector groups, of which 8 represented known or characterized effectors, while the remaining 13 had not previously been described in *Aeromonas*. We experimentally validated our *in silico* findings by assessing the cytotoxicity of representative effectors in *Saccharomyces cerevisiae* BY4741, with 15 out of 21 assayed proteins eliciting a cytotoxic effect in yeast. The results of this study demonstrate the utility of our approach, combining a novel *in silico* search method with *in vivo* experimental validation, and will be useful in future research aimed at identifying and authenticating bacterial effector proteins from other genera.

## Introduction

*Aeromonas* spp. are Gram-negative γ-proteobacteria, many of which are of increasing clinical significance as emergent human pathogens [1,2]. *Aeromonads* are present in diverse habitats and interact with a variety of organisms, such as leeches, zebrafish, vultures, and humans, participating in both mutualistic and pathogenic symbiotic interactions with their hosts [3–8]. *Aeromonas* species cause diseases in a variety of animals, and they are particularly associated with fish diseases caused by the psychrophilic species *A. salmonicida* as well as by mesophilic ones, such as *A. hydrophila*, *A. sobria* and *A. veronii* [5,9]. In humans, *Aeromonas* strains are identified as causative agents of traveler’s diarrhea and septicemia, as well as other severe infections such as necrotizing fasciitis [1,10].

Numerous important *Aeromonas* virulence factors have been identified, including aerolysin, exotoxins, and type three secretion systems (T3SSs) [11]. The T3SS is an important and one of the best-studied bacterial virulence factors [12–14], and the diversity of niches inhabited by *Aeromonas* spp. may be due in part to many strains possessing one or more T3SSs [15–21]. The T3SS is a molecular syringe that transfers effectors with a range of biochemical activities into target eukaryotic cells [22]. In addition, T3SSs are frequently associated with horizontal gene transfer (HGT) events and are commonly observed within pathogenicity islands [23–25]. Thus, the importance of T3SSs in bacterial-eukaryotic interactions and their frequent horizontal transfer makes them important players in niche adaptation processes [26].

Next-generation sequencing has driven remarkable advances in microbiology, both in fundamental research and clinical diagnostics [27,28]. A major objective in the acquisition and analysis of whole genome sequence data is to better predict the pathogenic potential of bacterial strains by identifying genetic determinants of virulence. The core components of the T3SS apparatus are homologous to the bacterial flagellum, and their sequences are sufficiently similar that cross identifications occur during homology searches. In contrast, T3SS effectors are less conserved and often display uncharacterized domains, which also leads to difficulties during homology searches. An example of these challenges are two effectors, AexT and AexU, in *Aeromonas*, which share one domain with a second domain being completely novel [29,30]. Databases such as the Pathosystems Resource Integration Center (PATRIC) [31], VICTORS [32], EffectiveDB [33], and the Virulence Factor DataBase (VFDB) [34] contain T3SS effector sequences with different levels of validation. One problem encountered when attempting to identify T3SS components through homology searches is that many proteins and protein domains encoded in bacterial genomes are homologs of T3SS components that are not part of a T3SS system, but rather encode other cellular components such as bacterial flagellar components [35], F-ATPase subunits [36], and type IV pili [37,38]. In this study, we describe an *in silico* approach that solves this problem through the use a negative control comprised of all protein sequences from *Vibrio fisherii* ES114 and *Escherichia coli* K12. Since these genomes are known to not encode a T3SS, their protein sequences can be used as non-T3SS references.

In this study, we report the distribution and cytotoxicity of 21 T3SS effector families within 105 *Aeromonas* genomes and describe the pipeline developed to identify them. The identified proteins were evaluated as potential T3SS effectors by expressing representative proteins from each family in the yeast *Saccharomyces cerevisiae* strain BY4741 and assessing their cytotoxicity. Out of 21 assayed effectors, 13 are newly described and 15 exhibited cytotoxicity under the conditions assayed. Our findings extend the knowledge of the breadth and distribution of T3SS effectors in *Aeromonas* strains, and our combined use of a bioinformatics pipeline followed by verification through heterologous expression in yeast provides a template for studies in other bacterial genera.

## Materials and Methods

### Sequences

All annotated protein coding gene sequences from the 105 *Aeromonas* strains were assessed in this study. From the 105 evaluated *Aeromonas* genomes, 40 genomes were sequenced during this study (S1 Table), 35 were previously published by our group [21,39,40], and 30 were obtained from public databases (S2 Table). The genomes were sequenced, assembled and annotated as described in Colston et al. 2014. Briefly, libraries were prepared using NexteraXT and sequenced on an Illumina MiSeq at the Microbial Analysis, Resources and Services Facility of the University of Connecticut. The reads were trimmed and assembled using CLC Genomics Workbench (Qiagen). Gene predictions and product annotations was performed using RAST [41]. The data is available under BioProject PRJNA391781 and SRA accession numbers SRS2335044-SRS2335083.

### Homology clustering

Protein sequences from the 105 *Aeromonas* genomes were clustered into homologous groups using the OrthoMCL algorithm [42] as implemented in the Get_Homologues software package [43]. A total of 25,518 homologous groups were assembled, 2,755 of which are present in at least 90% of surveyed genomes. The genes of this extended core comprise approximately 65% of the individual *Aeromonas* genomes.

### Identification of T3SS related proteins in *Aeromonas* spp. genomes

Reference amino acid sequences of T3SS apparatus and effector proteins were downloaded from the VFDB in April 2015. All *Aeromonas* spp. protein families were compared against the VFDB reference sequences using the HMMER suite [44]. HMMER profiles were generated for all identified homologous groups of *Aeromonas* proteins, whose alignments were performed using MAFFT [45]. Protein families without a positive VFDB hit (HMMER default parameters) were dismissed from further investigation. Proteomes of *V. fisherii* ES114 [46] and *E. coli* K12 [47,48] were used as negative controls as they are known to not to possess T3SSs. We reasoned that by removing homologous groups matching equally well, or better, to the negative controls as to the VFDB sequences that the inclusion of distant T3SS homologs, such as F-ATPase or flagellar components, would be avoided. All protein sequences from groups representing positive hits against the VFDB were used as queries in BLAST searches against the two different data sets, the VFDB reference sequences and the negative control. A t-test was performed on homologous groups that had at least three positive hits against both the VFDB and the negative control to evaluate if the sequences were significantly more similar to those in the VFDB compared to the negative control. In cases where the homologous group had less than three hits against one of the datasets, we assessed if the lowest alignment score from the VFDB was at least 1.5-fold greater than that observed from the negative control.

Gene families that were predicted to be related to the T3SS were manually curated and divided into separate clusters or merged into one as necessary. All putative effector families were manually identified from the set of 127 T3SS-related groups. The correlations between effector occurrences (S1 Fig) were assessed using Pearson’s correlation tests.

### Reference phylogeny

Multiple sequence alignments were generated for all 5,693 single copy gene families using MAFFT, and quality control and edits automatically performed by GUIDANCE [49]. Pairwise maximum likelihood distance matrices were calculated for gene families present in at least 10 genomes using Tree-Puzzle [50]. Pearson’s correlation tests between all gene family combinations were performed, and the significance of each correlation was assessed through Mantel tests and p-values corrected for multiple testing using Benjamini-Hochberg’s False Discovery Rate (FDR) correction. The correlation-weighted network (and) was submitted to the Markov Clustering Algorithm (MCL) with 5.5 inflation parameter [51,52]. The unpartitioned concatenation of the 1,678 gene families present in the largest cluster was submitted to RAxML [53] for phylogenetic reconstruction using the GTR+GAMMA substitution model, and support was assessed using the aLRT SH-like method [54].

### Gene phylogenies and reconciliations

Nucleotide sequences from all putative effector gene families were aligned using MAFFT, and phylogenetic trees were generated using RAxML (GTR+GAMMA+I model). The obtained trees were refined using the TreeFixDTL tool [55] and reconciled to the species tree using RangerDTL [56]. S2 Fig shows cladograms of putative effector phylogenies after applying the TreeFixDTL tool and rooting the gene trees through RangerDTL.

### Addition of non-*Aeromonas* representatives

Amino acid sequences from the putative T3SS effectors were used to query against GenBank to identify non-*Aeromonas* homologs. The top non-*Aeromonas* hit was downloaded if the aligned region spanned through at least 60% of the query sequence and had an identity of at least 30%. Once a non-*Aeromonas* hit fulfilling the requirements was identified, it was downloaded together with all other non-*Aeromonas* hits with bitscores of at least 85% of the top hit.

### Strains and growth conditions

The bacterial and yeast strains and plasmids generated in this study are listed in S1 Table. The *E. coli* strain DH5α λ-*pir* was used to clone the plasmids and was cultured in LB broth or on agar solidified plates containing 100 µg/ml ampicillin as needed. The yeast strain *S. cerevisiae* BY4741 was cultured using yeast extract-peptone-dextrose (YPD) medium for non-selective growth[57], and selective growth was performed using synthetic defined medium lacking histidine (SDM-His) and containing either 2% glucose (SDM-His-Glu), or 2% galactose and 1% raffinose (SDM-His-Gal). Some strains were additionally evaluated on SDM-His-Gal medium supplemented with 7 mM caffeine or 0.5 M NaCl.

### Strain and plasmid construction

All primers used in the present study are listed in Table S4. Putative T3SS effector genes were PCR amplified from *Aeromonas* genomic DNA using Phusion DNA polymerase (New England Biolabs; NEB) and appropriate primers. Primers were tailed to allow subsequent Gibson Assembly (NEB) cloning of amplicons at the *Eco*RI and *Xho*I sites of the shuttle vector pGREG533 (Euroscarf), a plasmid which allows for galactose inducible expression of cloned genes from the GAL1 promoter. Amplicons were purified using a Wizard SV Gel and PCR Clean up System kit (Promega) and *Eco*RI/*Xho*I digested pGREG533 was gel purified using a Qiaex II kit (Qiagen). The effector genes were cloned in-frame with the N-terminal 7× hemagglutinin tag (7-HA) in pGREG533. Constructs with inserts of the correct size were initially screened for by PCR and were subsequently sequenced and transformed into *S. cerevisiae* strain BY4741. Yeast were transformed by mixing 0.1 µg of plasmid, 0.1 mg of salmon sperm carrier DNA (Invitrogen) and 0.1 ml of yeast cells resuspended in sterile 1× TE/LiAC buffer (0.1 M Tris-HCl, 10 mM EDTA, and 0.1 M LiAc, pH 7.5). After adding 0.6 ml of sterile 1× TE/LiAc buffer containing 40% polyethylene glycol 4000, the cells were incubated at 30°C for 30 min with shaking (200 rpm). Next, the suspensions were mixed with 70 µl of dimethyl sulfoxide (DMSO), heat shocked at 42°C for ≥ 15 min, and then were centrifuged and resuspended in sterile water. Cell dilutions were plated on SDM-His-Glu plates and incubated for two days at 30°C to obtain transformants.

### Yeast growth inhibition assay

For each strain, 3 ml of SDM-His-Glu broth was inoculated with a single yeast colony and incubated at 30°C overnight with shaking. After centrifugation at 800 × *g* for 5 min, the supernatant was decanted and the cell pellet resuspended in sterile water. This process was repeated to remove residual medium, after which the cells were diluted to an OD_600_ of 1.0 and four 10-fold serial dilutions were made. A 10 μl aliquot of each dilution was spotted onto SDM-His-Glu plates or SDM-His-Gal. Yeast strains presenting partial (*aopP*, *pteH*, *pteJ* and *pteL*) or no growth inhibition phenotypes (*aopH*, *aopO*, *pteD*, *pteD.1*, *pteE* and *pteK*) on SDM-His-Gal plates were further assessed on SDM-His-Gal plates containing 7 mM caffeine or 0.5 M NaCl. The plates were incubated at 30°C for 2-3 days and were photographed with a Nikon D80 camera. Candidate effectors were cloned into pGREG533 in a galactose inducible manner from the GAL1 promoter. Plasmids expressing each of the 21 previously identified or putative effector proteins were separately introduced into the yeast strain *S. cerevisiae* BY4741. 10-fold serial dilutions of cells were spotted onto agar-solidified plates that repressed (SDM-His-Glu) or promoted (SDM-His-Gal) the expression of the protein of interest.

### Western blot analysis

The expression of putative T3SS effectors in yeast strains presenting no growth inhibition phenotypes under any of the assayed conditions (*aopH*, *aopO*, *pteD*, *pteD.1*, *pteE* and *pteK*) was assessed by western blotting. The strains were inoculated from plates into SDM-His-Glu broth and grown overnight at 30°C with shaking (200 rpm). One milliliter of each culture was pelleted and resuspended in 300 µl of lysis buffer and added to a bead beating tube containing 300 mg of 0.5 mm zirconia/silica beads (cat# 11079105z, BioSpec Products) (50 mM Tris-HCL, 1% DMSO, 100 mM NaCl, 1 mM EDTA, and 1 mM PMSF, pH 8.0). The samples were homogenized 2× at 2000 rpm for 20 seconds and 4× at 4000 rpm for 10 seconds using a Qiagen Powerlyzer 24, with the samples placed on ice between each round of bead beating. The beads were pelleted by centrifuging the samples at 16,000 × *g* for 1 minute, after which supernatants were mixed with an equal volume of Laemmli buffer and were heated for 5 min at 100°C. The protein extracts were separated by SDS-PAGE on a 4-20% Mini Protean TGX gel (Bio-Rad) and then transferred to a PVDF membrane. The N-terminal 7-HA-tagged proteins were probed for using a mouse monoclonal anti-HA antibody followed by an HRP-conjugated goat anti-mouse IgG H&L HRP secondary antibody (ab49969 and ab205719, respectively; Abcam). The blots were developed using an ECL Plus Western Blotting Substrate kit (Pierce) and imaged using a FluorChem HD2 (ProteinSimple).

## Results and discussion

### Reference phylogeny

A phylogenomic study of 56 high-quality genomes from *Aeromonas* strains was published together with an MLSA tree of 16 housekeeping genes from the same genomes [40]. This study identified incongruities between gene, MLSA, and core genome trees [58], which is expected since different sets of genes were used to construct each tree. In our current study, we expanded on this set with an additional 49 *Aeromonas* genomes. To produce a reference phylogeny that reflects a significant share of *Aeromonas* genes and minimize conflicting evolutionary signals among genes, we constructed a phylogenomic tree using 1,678 genes with compatible evolutionary histories. Building phylogenies from concatenated genes is a widely used approach to reconstruct the representative history of a genome. However, the concatenation of genes with different trajectories can lead to unresolved trees and/or a phylogeny that represents neither the history of the organism nor that of the individual genes used in the concatenation [59–62]. One approach used to obtain better phylogenies is to group gene families with compatible evolutionary histories using tree distance metrics [63,64]. Therefore, to assess the similarity between the evolutionary histories of different genes we performed pairwise Pearson’s correlation tests between Maximum Likelihood distance matrices of all gene families present in at least 10 genomes. It has been discussed that metrics accounting for both topology and branch length do not perform significantly better than metrics accounting only for the latter [64]. For comparison, we also measured the distances between gene families using the Fitch-Margoliash criterion [65], i.e., in calculating the sum of squares of the differences in distance between two matrices, the square of each difference was multiplied by the inverse of the distance, thereby increasing emphasis on the difference in distance between more similar sequences. The two approaches used to compare distance matrices showed a strong correlation (). Significant correlation coefficients, were submitted to a Markov Clustering (MCL) process [52], and a weighted network of connected gene families were constructed as discussed by Dongen and Abreu-Goodger (2012). From the 168 gene family clusters obtained through MCL clustering, we considered the largest cluster, which contains 1,678 gene families, to be the primary phylogenetic signal among *Aeromonas* strains. This gene family cluster comprises more than twice the number of gene families present in the second largest cluster, which contains 738 gene families. By using the largest gene cluster, containing gene families with different functions and genomic locations but with compatible evolutionary histories, we ensured that our phylogeny is representative of the evolutionary history of the genomes.

On average, each *Aeromonas* genome has homologs from 1,637 out of the 1,678 families used in the reference phylogeny (), representing 39% of the average number of coding sequences in the *Aeromonas* species assayed. The resulting phylogenomic tree (Fig 1) is highly supported, with all but three branches displaying an aLRT SH-like support [54] larger than 0.90. The concatenated gene tree presented in this study is largely consistent with the previously published *Aeromonas* spp. core-genome phylogeny [40], with the only differences being the deeper branching of *A. salmonicida* strains in relation to the common ancestor of *A. hydrophila* and *A. veronii*, as well as *A. allosaccharophila* being a sister group of *A. veronii*.

**Fig 1.**
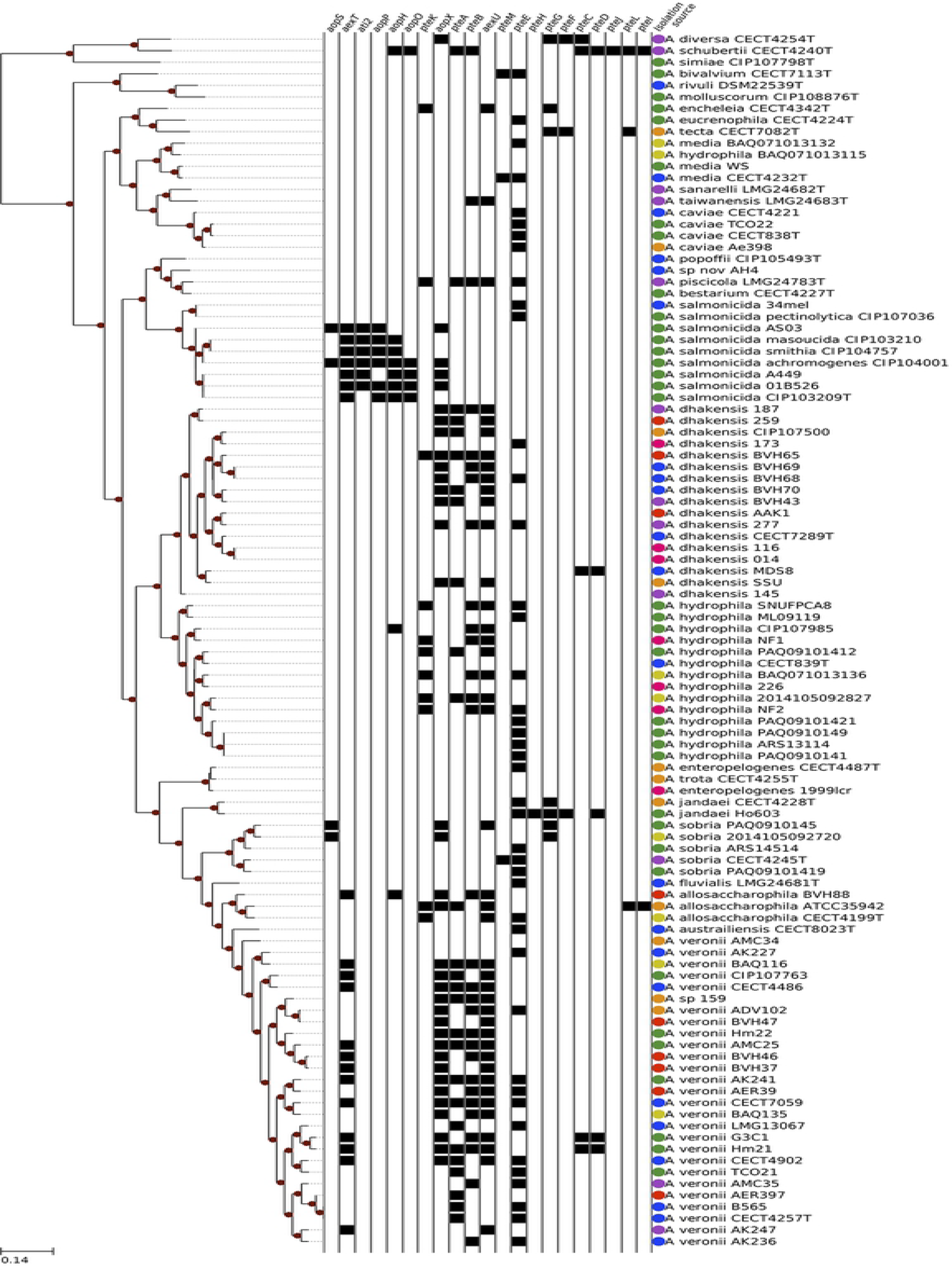
Phylogeny of 105 *Aeromonas* spp. genomes. Tree branches with sh-like support of at least 90% are highlighted with red circles. The binary heat map represents the presence/absence of putative effectors in *Aeromonas* spp. isolates. The isolation source colors represent: green, veterinary; yellow, sick veterinary; blue, environment; orange, feces; red, blood; purple, wound; and pink, human.

### *In silico* identification of putative T3SS effectors

For the identification of putative effectors, initially all coding sequences from sampled genomes were compared to each other to cluster homolog genes. Hidden Markov Model (HMM) profiles generated from each of the 25,518 identified *Aeromonas* spp. homolog gene families were queried against T3SS sequences from the VFDB, which yielded 633 positive hits. Although some of these positive hits were clearly not T3SS related they displayed alignments with e-values as low as due to a common origin of other systems shared with T3SS components. Protein sequences from *Vibrio fisherii* ES114 [46] and *Escherichia coli* K12 [48] were used as negative controls for bona fide T3SS sequences, as both species are known to not have a T3SS. The combination of BLAST searches of sequences from the 633 initial candidates against both VFDB and the negative control genomes identified 127 gene families significantly more similar to T3SS sequences. Through subsequent manual curation, we assessed the domains present in each putative family and their genomic contexts (*e.g.*, whether they are adjacent to chaperones) and classified 21 gene families as encoding likely effectors (Table 1).

**Table 1.** Relevant characteristics of putative T3SS effector groups. Number of proteins gives the total number of protein encoding genes identified across all assayed genomes for a given group.

| Effector | # of Proteins<br>(# of T<sup>s</sup>) | Length in bp<br>(max, median,<br>min)<sup>\$\$</sup> | Pfam<br>Domains <sup>\$\$\$</sup> | Domain Descriptions <sup>\$\$\$</sup> | GOterm IDs |
| --- | --- | --- | --- | --- | --- |
| **AexT** | 20 (5-5) | 1429, 1353, 1287 | PF03496;<br>PF03545 | ADP-ribosyltransferase<br>exoenzyme; Yersinia virulence<br>determinant (YopE) | GO:0005576,<br>GO:0009405 |
| **AexU** | 42 (15-15) | 1545, 1537.5, 768 | PF03545 | Yersinia virulence determinant (YopE) | N/A |
| **AopX** | 36 (14-14) | 972, 951, 948 | N/A | N/A | N/A |
| **AopO** | 5 (0-1) | 2188, 2187, 2187 | PF09632;<br>PF00069 | Rac1-binding domain; Protein kinase domain | GO:0004672,<br>GO:0005524,<br>GO:0006468 |
| **AopP** | 6 (0-0) | 897 | PF03421 | YopJ Serine/Threonine acetyltransferase | N/A |
| **Ati2** | 6 (0-0) | 1488, 1488, 1242 | PF03372 | Endonuclease/Exonuclease/ phosphatase family | N/A |
| **AopH** | 9 (0-3) | 1392, 1206, 1152 | PF00102;<br>PF09013 | Protein-tyrosine phosphatase; YopH, N-terminal | GO:0004725,<br>GO:0006470,<br>GO:0004725,<br>GO:0006470,<br>GO:0009405 |
| **AopS** | 5 (0-1) | 1155, 1155, 1149 | PF02661 | Fic/DOC family | N/A |
| **PteA** | 27 (0-13.5) | 2112, 2103, 1944 | N/A | N/A | N/A |
| **PteB** | 30 (4-18) | 951, 903, 492 | N/A | N/A | N/A |
| **PteC** | 5 (0-2) | 1044, 1044, 999 | N/A | N/A | N/A |
| **PteD** | 4 (0-2) | 1089, 1083, 1083 | N/A | N/A | N/A |
| **PteD.1** | 1 (0-0) | 1032 | N/A | N/A | N/A |
| **PteE** | 51 (0-31.9) | 1443, 1128, 1083 | PF13776;<br>PF02661 | Domain of unknown function (DUF4172); Fic/DOC family | N/A |
| **PteF** | 3 (2-0) | 1059, 969, 963 | PF03543 | Yersinia/Haemophilus virulence surface antigen | GO:0004197,<br>GO:0009405 |
| **PteG** | 7 (0-2.79) | 2010, 1872, 1833 | PF01734 | Patatin-like phospholipase | GO:0006629 |
| **PteH** | 1 (0-0) | 1383 | PF01734 | Patatin-like phospholipase | GO:0006629 |
| **PteI** | 2 (1-1) | 1134, 1132.5,<br>1131 | PF03497 | Anthrax toxin LF subunit | GO:0005576,<br>GO:0008294,<br>GO:0009405 |
| **PteJ** | 1 (0-0) | 708 | PF03536 | Salmonella virulence-associated 28kDa protein | N/A |
| **PteK** | 11 (0-4.86) | 1422 | PF09013 | YopH, N-terminal | GO:0004725,<br>GO:0006470,<br>GO:0009405 |
| **PteL** | 3 (2-2) | 1452, 1452, 1446 | N/A | N/A | N/A |
| **PteM** | 2 (1-1) | 678, 615, 552 | PF03496 | ADP-ribosyltransferase<br>exoenzyme | GO:0005576,<br>GO:0009405 |
| **PteM.1** | 1 (0-0) | 1098 | PF03496 | ADP-ribosyltransferase<br>exoenzyme | GO:0005576,<br>GO:0009405 |

\$. Duplication, transfer, and loss events were estimated using Ranger-DTL [56], see material and methods for details. No duplication was predicted within the putative effectors evolution. The number of transfers with 100% confidence and the mean number of transfer events is given in parentheses.
\$\$: In instances where the max, median and min nucleotide length of genes was identical a single value is given.
\$\$\$: Domains present in an effector group were identified using the Pfam database, and the PFAM associated Go term ID is given.

Eight of the 21 likely effectors had been previously identified in *Aeromonas* species, including *aexT* [17,19,66], *aexU* [67], *aopP* [68,69], *aopH* and *aopO* [70], *ati2* [71], and *aopX* and *aopS*, which were initially identified in *A. salmonicida* as pseudogenes [72]. We also identified *aopN*, but did not consider it further in our study as it was shown in *Bordetella bronchiseptica* to have a dual role in controlling the secretion of translocator proteins and suppressing host immunity but was not cytotoxic [73].

The remaining 13 likely *Aeromonas* T3SS effectors had not been studied in any detail and were designated as <u>p</u>utative <u>T</u>3SS <u>e</u>ffectors, which included *pteA*, *pteB*, *pteC*, *pteD*, *pteD.1, pteE*, *pteF*, *pteG*, *pteH*, *pteI*, *pteJ*, *pteK*, and *pteL. pteD.1* was combined with the pteD family despite being originally classified into distinct homolog groups since they share significant sequence similarity, as assessed through PRSS [74] using the PAM250 scoring matrix. The comparison between PteD.1 from A. jandaei Ho603 and PteD from A. sp. MDS8 sequences resulted in a Z-score of 65.7. Besides significant sequence similarity, homologs from both pteD and pteD.1 are likely related to the same chaperone family, whose members were automatically grouped within a single homolog protein cluster. The thirteen newly described putative T3SS effectors and the 8 previously described effectors were submitted to InterProScan [75] for domain identification (Table 1), and were subsequently assessed for cytotoxic effects through expression in *S. cerevisiae* strain BY4741.

### Screening of putative T3SS effector proteins in yeast

To assess the cytotoxicity of putative T3SS effectors identified in *Aeromonas* spp. genomes, we expressed representative proteins from each putative group in the yeast strain *S. cerevisiae* BY4741 and monitored for growth inhibition (Table 1). Although eight of the identified *Aeromonas* effector families have previously been identified or studied, to the best of our knowledge, none have been assessed for causing cytotoxicity or growth inhibition in yeast. Expression of bacterial toxins in yeast is a common means of assessing the deleterious impact of these proteins on eukaryotic host cell processes [76–78]. The serial dilutions allow one to assess better how severe the cytotoxicity of an effector is. While all of the strains grew on medium containing glucose (Fig 2), those expressing *pteA* and *pteF* were somewhat inhibited for growth under non-inducing conditions, yielding colonies with reduced size relative to cells carrying the pGREG533 plasmid alone. Presumably, these effectors are so cytotoxic that even uninduced, basal protein expression is sufficient to inhibit growth. Interestingly, yeast cells transformed with *pteF* yielded colonies with different sizes, possibly due to the acquisition of suppressor mutations in the larger colonies.

**Fig 2.**
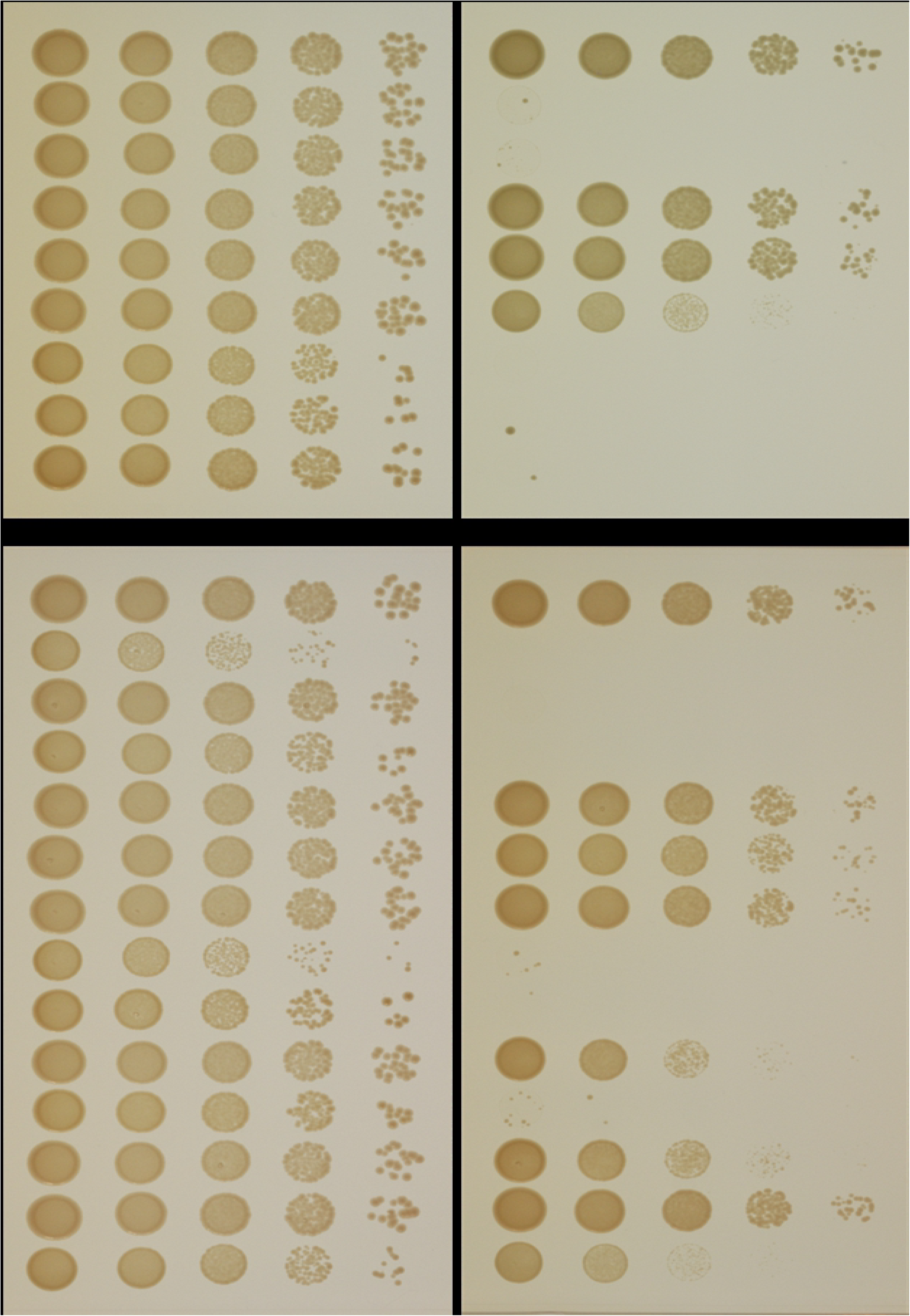
Yeast growth inhibition assay. Strains carrying putative T3SS effectors cloned into pGREG533 were cultured overnight in SDM-His-Glu, washed and 10-fold serially diluted. Aliquots from each dilution (10 µl) were spotted onto SDM-His-Glu and SDM-His-Gal plates. A strain containing pGREG533 was used as a negative control and showed no growth inhibition. SDM-His-Glu and SDM-His-Gal plates were incubated at 30°C for 2-3 days. T3SS effectors previously identified or biochemically characterized are presented in panel (A), whereas those identified in the present study are shown in panel (B).

Yeast strains expressing 15 of the 21 assayed proteins exhibited a growth phenotype when plated on SDM-His-Gal inducing medium (Fig 2). Strains expressing the proteins encoded by *aexT*, *aexU*, *aopS*, *aopX*, *ati2*, *pteA*, *pteB*, *pteC*, *pteF*, *pteG* and *pteI* showed little or no growth on the 10^0^ dilution (Fig 2), indicating that these effectors are very cytotoxic. In addition, strains expressing *aopP*, *pteH*, *pteJ*, and *pteL* exhibited reduced colony sizes with either no or little growth on the 10^−4^ dilution, suggesting that they are less cytotoxic. Strains expressing *aopH*, *aopO*, *aopD*, *aopD.1*, *pteE*, or *pteK* did not exhibit a growth inhibition phenotype compared to the control strain harboring pGREG533.

If the cellular process targeted by a bacterial effector does not typically limit yeast growth, the presence of stressors (*e.g.*, elevated salt or caffeine) can promote the inhibition phenotype to be observed [79]. The addition of NaCl to SDM-His-Gal medium inhibited the growth of strains expressing *aopP* and *pteJ* (slight growth on the 10^0^ dilution) compared to that observed on unsupplemented SDM-His-Gal medium (growth on the 10^4^ dilution) (Fig 3). On SDM-His-Gal plates containing caffeine, no further reduction in growth was observed for the *aopP*-expressing strain, whereas that of *pteJ*-expressing strain was observed on the 10^0^ and 10^−1^ dilutions (Fig 3). In addition, the strains expressing *pteH* and *pteL* produced small colonies at the 10^−3^ and 10^−2^ dilutions, respectively, when grown on SDM-His-Gal medium containing NaCl compared to that observed on SDM-His-Gal medium alone (growth on the 10^−3^ and 10^−2^ dilutions, respectively) (Fig 3). On plates containing caffeine, the strains expressing *pteH* and *pteL* grew on the 10^−3^ and 10^−1^ dilutions, respectively (Fig 3). For the constructs that did not produce a growth inhibition phenotype (*aopH*, *aopO*, *pteD*, *pteD.1*, *pteE*, and *pteK*), we assessed whether bacterial proteins of the expected fusion protein size were expressed by western blot and all of proteins were expressed at the predicted size except PteD for which no product was detected (S3 Fig).

**Fig 3.**
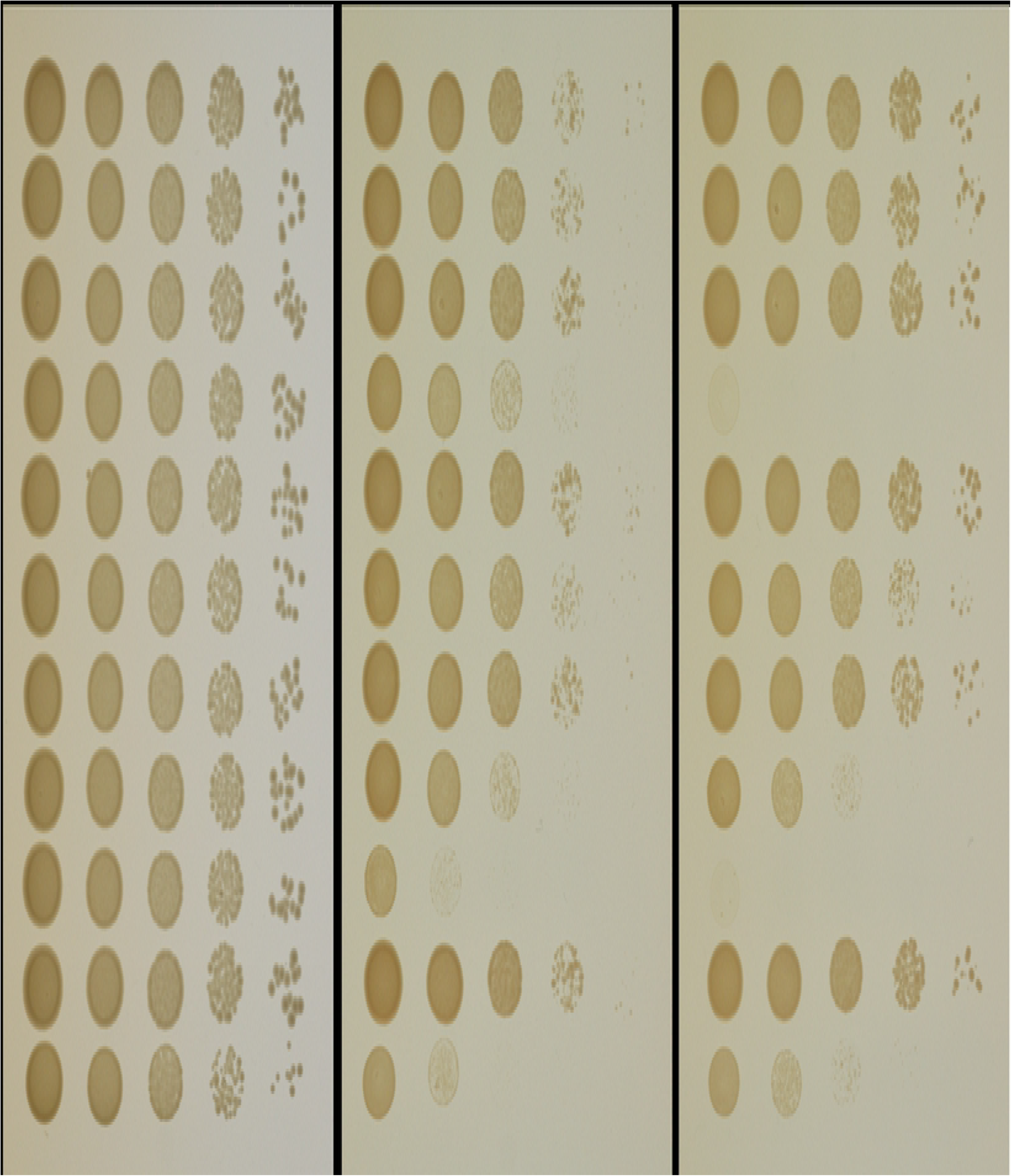
Yeast growth inhibition assay under stress conditions. Strains carrying putative T3SS effectors cloned into pGREG533 were grown overnight in SDM-His-Glu, washed and 10-fold serially diluted. Aliquots from each dilution (10 µl) were spotted onto SDM-His-Glu or on SDM-His-Gal plates containing either 0.5 M NaCl or 7 mM caffeine. The strain containing pGREG533 was used as a negative control and showed no growth inhibition. The plates were incubated at 30°C for 2-3 days.

Both AopH and PteK have similar domain structures (Table 1), and neither protein elicited a growth phenotype when expressed in yeast. The *Yersinia* homologs, YopH and YpkA, of these two *Aeromonas* proteins have been previously assayed in yeast for cytotoxicity [80]. Although YopH similarly did not cause a growth inhibition phenotype, YpkA strongly inhibit yeast growth. Several explanations could account for the lack of observed phenotypes in strains expressing AopH, AopO, PteD, PteD.1, PteE, and PteK, including: a lack of activity due to the adjoined 7-HA tag; the lack of a target protein for the effector in the yeast strain assayed; the effector does not produce a cytotoxic effect but interacts with host cells in another manner; the protein was not expressed at a high enough level; or the putative effector may not be a bacterial toxin.

These assays allowed the putative T3SS effectors identified in our bioinformatics analysis to be rapidly evaluated. Our findings showed that 15 out of the 21 tested proteins inhibited growth of *S. cerevisiae* BY4741, and that addition of NaCl, and to a lesser extent caffeine, to SDM-His-Gal plates increased the sensitivity of yeast cells to four of these likely effectors. Future assessments of the effectors with no observed phenotypes should focus on generating C-terminally tagged proteins, since the N-terminal tags generated in the current study could be the cause of aberrant activity or localization that may potentially mask a growth or cytotoxicity phenotype. Additionally, alternate yeast strains could be tested.

### Distribution and evolutionary history of putative effectors in *Aeromonas*

The scattered occurrence of the 21 predicted effectors described in this study throughout the *Aeromonas* phylogenomic tree is evidence of the impact of HGT during their evolution. Since only 15 out of the 21 putative effectors are present in at least four taxa, our HGT inferences using phylogenetic reconciliation are restricted to *aexT, aexU, aopX, aopO, aopP, ati2, aopH, aopS, pteA, pteB, pteC, pteD, pteE, pteG*, and *pteK*. In effectors present in less than four genomes, we only considered their presence/absence to infer gene transfer and loss events. The presence of a putative effector among Aeromonads has an origin in two possible scenarios: [1] vertical inheritance, or [2] HGT from a non-*Aeromonas* lineage. In a reconciliation scenario where no HGT is allowed among Aeromonads, 1,177 gene loss events and 105 gene duplication events are required to reconcile the histories of 15 putative effectors across *Aeromonas* phylogeny. In the reconciliation of the same 15 putative effectors allowing HGT and using default reconciliation penalties (loss = 1, duplication = 2, and transfer = 3), only 85 gene losses and 115 HGTs are required. Of the 115 predicted HGT events, 44 took place between terminal nodes of the tree. There is a significantly larger number of inferred HGTs between genomes from distinct species (30) than between members of the same species (14). Such small number of inferred within-species transfers may be due to a lack of resolution in the gene trees leading to inferred events with low confidence values. If a strong phylogenetic signal is absent from bipartitions present in the putative effector tree, our reconciliation approach assumes it to be equivalent to the genome phylogeny, since the gene tree bipartition does not strongly support the incongruence. Reconciliations also revealed a large variance of HGT events inferred among T3SS effector gene families, ranging from 0 to 32 horizontal transfers within *Aeromonas* spp.

In order to evaluate the gene exchange between *Aeromonas* spp. and other lineages, we recruited non-*Aeromonas* homologs of the 15 putative effectors present in more than four *Aeromonas* genomes. The inclusion of non-*Aeromonas* homologs revealed that *Aeromonas* T3SS effectors are not frequently shared outside the genus boundaries. In all extended putative effector trees, *Aeromonas* homologs grouped into clans [81], excluding homologs from other taxa to separate clades. This result suggests that each of the putative effectors has a single origin within *Aeromonas* strains, i.e., they entered the genus only once, either from its common ancestor or through a single HGT event. The disparity between within-genus and between-genera transfers is well described in the literature [20,82–84]. The high number of inferred gene transfers within *Aeromonas* genomes is evidence of the interchangeability of T3SS effectors within the genus, while the paucity of HGT events between genera may either reflect decreasing transfer rate with increasing evolutionary distance [83–85] or the high degree of specificity between effectors and the secretion system apparatus. We recognize the potential for undetected transfers with unsampled lineages not represented in public databases, although one would expect to observe at least some number of paraphyletic aeromonad clades if between-genera exchanges involving *Aeromonas* were common.

Five putative effector families exhibited more than ten HGT events within *Aeromonas* spp. according to phylogenetic reconciliations. The most exchanged effector is *pteE*, with 32 inferred transfers within the genus. The reconciliation analysis inferred 18 HGT events during the evolution of *pteB* in *Aeromonas*. Two of the identified effectors, *aopP* and *ati2*, did not require horizontal transfers during its reconciliations with the genome phylogeny. Both effector families are exclusively present in *A. salmonicida* genomes, although *aopP* is absent from *A. salmonicida* A449, and *ati2* is absent from *A. salmonicida* CIP103209T, and their closest non-*Aeromonas* homologs are present in *Yersinia enterocolitica* and *Vibrio harveyi*, respectively. The genomes of *A. salmonicida* strains are very similar to each other, which is reflected in the short branch lengths present in the genome phylogeny (Fig 1). Thus, we would not expect a significant number of different HGT events among *A. salmonicida* strains. We hypothesize that an *A. salmonicida* common ancestor acquired these effector genes through horizontal transfer, likely from *Yersinia* spp. or *Vibrio* spp., which were then vertically inherited by most of the descendants.

The great variation in effector occurrence among *Aeromonas* strains, from zero to nine effectors per genome, may reflect the diversity of lifestyles observed in this genus. For example, *A. schubertii* possesses nine likely effectors in its genome, more than any other assessed strain, and it is also among the earliest branching aeromonads (Fig 1). We were unable to determine if this large number of predicted T3SS effectors is related to the niche *A. schubertii* occupies (cultured from a clinical forehead abscess), since no other sampled strain was isolated from a similar source. On the other side of the spectrum, 19 strains have no putative effector, and they are found across all clades of the *Aeromonas* phylogeny. Among putative effectors, some exhibit a wide distribution (*pteA* and *pteB*), and others have a very restricted distribution (*pteJ*, *pteI*, and *pteL*).

The majority of the putative effectors display significant co-occurrence with other putative effectors in *Aeromonas* genomes (). One cluster of five putative effectors displays strong co-occurrence (*aexT*, *aopH*, *aopO*, *aopP* and *ati2*) (S1 Fig), and their occurrence is most prevalent in the *A. salmonicida* branch (Fig 1). Interestingly, one effector in this co-occurring cluster, *aexT*, is also present in the *A. veronii* strains. Another co-occurring cluster comprises five putative effectors (*aexU*, *aopX*, *pteA*, *pteB* and *pteK*) (S1 Fig), and their occurrence is mainly related to two different clades of the phylogeny, being identified among members of the *A. hydrophila*, *A. dhakensis*, and *A. veronii* clades. Despite the co-occurring clusters, we are unable to identify links between presence/absence of putative effectors and isolation sources of their respective genomes. The resemblance of phylogenetic signal observed in co-occurrence of putative effectors is probably due to the higher within-species HGT frequency, as once the effector is acquired by a member of the species it is easily spread among closely related genomes [85,86]. Due to the low frequency of putative effectors in the remaining two clusters displayed in S1 Fig, we cannot reliably infer what led to their co-occurrence.

## Conclusions

In this study, we identified likely T3SS effectors present in the genomes of 105 *Aeromonas* strains and assessed their cytotoxicity in *S. cerevisiae*. The *in silico* identification of T3SS effector sequences has been considered to be a complex task given that their short sequences and shared homology with proteins associated with different cellular systems constitute a barrier to accurate analysis [87–89]. Our two-step comparisons against positive and negative data sets greatly reduced the number of false positives and resulted in the identification of 13 new likely *Aeromonas* effector families and eight that were previously described. The expression of these proteins in *S. cerevisiae* provided strong evidence for the cytotoxicity of most of the identified effectors.

Based on the *in silico* and *in vitro* evidence obtained in this study, we propose naming the 9 newly described effectors with validated cytotoxic activity with the prefix *“ate*”, for <u>*A*</u>*eromonas* <u>T</u>3SS <u>e</u>ffectors. Consequently, the newly described *Aeromonas* effector families should be referred as *ateA*, *ateB*, *ateC*, *ateF*, *ateG*, *ateH*, *ateI*, *ateJ*, and *ateL*, whereas *pteD*, *pteD.1 pteE* and *pteK* should keep the putative designation since no yeast growth inhibition phenotype was detected.

The high frequency of horizontal transfer events of effectors within *Aeromonas* is reflected in their scattered distribution throughout the phylogenomic tree of the genus and reconciliations of each effector gene tree with the phylogenomic tree. Members of the *Aeromonas* genus are known for promiscuous gene exchange [20,90], as are genes related to the T3SS [91,92]. A comparison of the number of DTL events necessary for gene tree reconciliations between scenarios where HGT was allowed or not provided strong evidence for the high exchange rate of putative T3SS effectors among aeromonads. The larger number of predicted inter-versus intra-species HGTs could be explained by vertical transmission being the dominant mode in any given species. In the context of our study we observed that sharing an isolation source had a smaller impact on effector distribution compared to the phylogenetic signal. This result could also be due to *Aeromonas* strains interacting with a wide range of eukaryotic hosts, with each strain requiring a different set of molecular tools. Again, this strong phylogenetic signal in T3SS effectors distribution reflects the importance of the vertical inheritance of effectors within closely related organisms.

Using a combination of bioinformatic and molecular approaches, we were able to identify nine new T3SS effectors that are toxic to yeast cells. Future studies focusing on the role of individual effectors in different animal models, e.g., fish, mice, wax worms, and leeches would be useful in further characterizing these effectors and perhaps identify specific niches that some are associated with. In addition, the bioinformatic approach that we described can be used to identify effectors in other genera and also be applied to other gene families.

## Acknowledgements

This work was supported by the USDA ARS agreement 58-1930-4-002 and grants from the National Science Foundation (NSF/MCB #1616514). The Illumina sequencing was performed at the Microbial Analysis, Research and Services facility of the University of Connecticut. LTR was supported by the Fundação de Amparo a Pesquisa do Estado de São Paulo (FAPESP) through awards 2012/17196-2 and 2014/23975-0.

## Supporting Information

**S1 Figure. Co-occurrence correlation heat map.** The heat map displays all significant co-occurrences () between putative effectors in *Aeromonas* spp. isolates. The assessed correlation scale varies between 0.8 and −0.4. The four observed clusters are composed of: [1], *aexT*, *aopH*, *aopO aopP*, *aopS* and *ati2*; [2] *aexU aopX*, *pteA*, *pteB*, and *pteK*; [3] *pteF*, *pteH*, and *pteG*; and [4] *pteC*, *pteD*, *pteI*, *pteJ*, and *pteL*.

**Figure S2. Phylogenies for the putative effectors.** The depicted root corresponds to a most parsimonious DTL reconciliation. Numbers give percent bootstrap support calculated from 1000 samples using RAxML (GTR+GAMMA+I model).

**S3 Figure. Western blot analysis of *S. cerevisiae* BY4741 lysates expressing the 7-HA epitope-tagged putative T3SS effectors *aopH*, *aopO*, *pteD*, *pteD.1 pteE* and *pteK*.** The molecular weights of the of the major bands corresponded to those of the indicated effectors as determined by comparison with Kaleidoscope prestained standard (BioRad).

S1 Table. Characteristics and accession numbers for the genomes first reported in this study.

S2 Table. List of the 105 *Aeromonas* strains used in this study. S3 Table. Strains and plasmids used in this study.

S4 Table. Primers used in this study.

